# Combining the CRISPR Activation and Interference Capabilities Using dCas9 and G-Quadruplex Structures

**DOI:** 10.1101/2024.11.19.624357

**Authors:** Mohammad Lutful Kabir, Sineth G. Kodikara, Mohammed Enamul Hoque, Sajad Shiekh, Janan Alfehaid, Soumitra Basu, Hamza Balci

**Affiliations:** Department of Chemistry and Biochemistry, Kent State University, Kent, OH 44242, USA; Department of Physics, Kent State University, Kent, OH 44242, USA; Department of Physics, College of Science, Northern Border University, Arar, Saudi Arabia; Department of Chemistry and Biochemistry, Texas State University, San Marcos, TX 78666, USA

**Keywords:** CRISPR, dCas9, G-quadruplex, Transcription Regulation, CRISPRi, CRISPRa

## Abstract

We demonstrate that both CRISPR interference and CRISPR activation can be achieved at RNA and protein levels by targeting the vicinity of a putative G-quadruplex forming sequence (PQS) in the *c-Myc* promoter with nuclease-dead Cas9 (dCas9). The achieved suppression and activation in Burkitt’s Lymphoma cell line and in *in vitro* studies are at or beyond those reported with alternative approaches. When the template strand (contains the PQS) was targeted with CRISPR-dCas9, the G-quadruplex was destabilized and *c-Myc* mRNA and protein levels increased by 2.1-fold and 1.6-fold, respectively, compared to controls in the absence of CRISPR-dCas9. Targeting individual sites in the non-template strand with CRISPR-dCas9 reduced both the *c-Myc* mRNA and protein levels (by 1.8-fold and 2.5-fold, respectively), while targeting two sites simultaneously further suppressed both the mRNA (by 3.6-fold) and protein (by 9.8-fold) levels. These were consistent with cell viability assays when single or dual sites in the non-template strand were targeted (1.7-fold and 4.7-fold reduction in viability, respectively). We also report extensive *in vitro* biophysical studies which are in quantitative agreement with these cellular studies and provide important mechanistic details about how the transcription is modulated via the interactions of RNA polymerase, CRISPR-dCas9, and the G-quadruplex.

## INTRODUCTION

Clustered regularly interspaced short palindromic repeats (CRISPR) and CRISPR-associated (Cas) proteins are found in bacteria and archaea, functioning as an RNA-driven adaptive immune system against invading bacteriophages (1, 2). By using a modified version of Cas9, called nuclease-deficient Cas9 (dCas9), this system can be adapted to target genomic DNA without cutting it (3), as demonstrated by mutating the HNH and RuvC nuclease domains of the *S. pyogenes* Cas9 (3, 4). The dCas9 and a guide RNA (gRNA) complex has been utilized to regulate transcription in a sequence specific manner. To activate or boost transcription, dCas9 has been fused to a transcriptional activator domain, which helps recruit RNA polymerase (RNAP) (5–8). CRISPR-dCas9-mediated transcription activation (CRISPRa) is simple, highly specific, programmable, and more versatile compared to traditional gene expression modulation methods that rely on modifying genes and promoters (5, 9), as it primarily functions via recruitment of various effector proteins (10, 11). In *E. coli*, CRISPR interference (CRISPRi) has demonstrated dCas9’s ability to repress genes by blocking transcription elongation or preventing transcription initiation by interfering with transcription factor binding (3, 12–16). In this study, we demonstrate transcription regulation via a synergistic use of CRISPR-dCas9 and G-quadruplex (GQ) structures to achieve both CRISPRa and CRISPRi within the same system (17).

Genome-wide computational studies and high-throughput sequencing have identified several hundred thousand intramolecular putative GQ forming sequences (PQS) in the human genome (18). Telomeres and promoters, especially the immediate vicinity of transcription start site (TSS), are rich in PQS, suggesting a role in transcription level gene expression regulation (19–21). About 50% of human genes contain a PQS within 1,000 nucleotides (nts) upstream of TSS (22). PQS are more prevalent in promoters of oncogenes and regulatory genes, such as transcription factors, compared to housekeeping genes (23) and have been demonstrated to be involved in transcription regulation (17–21). Therefore, being able to regulate transcription by targeting the PQS with dCas9 promises to be a widely applicable and sequence specific method of transcription regulation, unlike alternative methods (such as, small molecules or site directed mutagenesis) which lack sequence specificity or transience. A prominent PQS located in the promoter region of *c-Myc* gene is used as a model system in this study.

The c-Myc is an oncoprotein and a transcription factor that plays an essential role in cell proliferation and induction of apoptosis (24–26). Overexpression of *c-Myc* is associated with a significant number of human malignancies, including breast, colon, cervix, and small-cell lung cancers, osteosarcomas, glioblastomas, and myeloid leukemias (27, 28). The *c-Myc* transcription is under the complex control of multiple promoters. The nuclease hypersensitivity element III1 (NHE III1) in the proximal region of the *c-Myc* promoter (−142 to −115 base pairs) can form two highly stable GQs and controls 80–90% of the total transcriptional activity of this gene (29, 30). It was demonstrated that stabilizing this PQS with small molecules reduced transcription by ∼2-fold while destabilizing the GQ with site-specific sequence mutations enhanced transcription by ∼3-fold in a Burkitt’s lymphoma cell line (31). On the other hand, a recent study reported that transcription factors preferentially interact with the GQ structure in the MYC promoter, and the GQ serves as positive regulator of transcription (32), opening the role of this GQ structure to further investigations.

In the absence of a PQS, dCas9 blocks RNAP from *E. coli* and bacteriophages SP6, T3, and T7 to different extents (12). Even though both dCas9 and GQs can independently block RNAP progression, dCas9 alone may also stabilize or destabilize the GQs (depending on whether the G-rich or C-rich strand is targeted) (33); thus, consolidating the two effects and potentially providing a broader range for transcription regulation. When targeting the PQS or the complementary sequence, dCas9 may even promote RNAP progression by destabilizing the GQ or further suppress it by stabilizing the GQ, and itself act as an additional blockade. Judiciously tuning these effects by properly selecting the dCas9 target site within the vicinity of PQS may enable up or down regulation of transcription. We present an example of these capabilities in this study.

We demonstrate that targeting the vicinity of *c-Myc* PQS with CRISPR-dCas9 provides levels of regulation that match or exceed those enabled with small molecules and sequence mutations. Furthermore, we demonstrate that whether the target sequence of dCas9 is in the template strand (TS) or non-template strand (NTS) of transcription is critical not only because of the resulting modulation in GQ stability, but also because of the different levels of blockade CRISPR-dCas9 presents for the RNA polymerase (RNAP) in these cases. Finally, we demonstrate that targeting two sites simultaneously with CRISPR-dCas9 enhances the range of regulation and enables more prominent levels of transcription suppression for *c-Myc*.

## METHODS

### Oligonucleotide preparation

All RNA and DNA oligonucleotides sequence information are reported in Tables S1 (33). The sequence of the 27 nt long PQS of the *c-Myc* promoter is follows: T<u>GGGG</u>A<u>GGG</u>T<u>GGGG</u>A<u>GGG</u>T<u>GGGG</u>AAGG, with the underlined five G-tracts shown to form two different GQ structures (34).

We used separate strands for CRISPR-RNA (crRNA) and transactivating CRISPR RNA (tracrRNA) in the CRISPR-dCas9 complex. Annealing these two strands resulted in the guide RNA (gRNA). The tracrRNA and all crRNA components used for the *c-Myc* system were *in vitro* transcribed in the lab. All DNA oligonucleotides (including those used as template for *in vitro* transcription) were purchased from either Integrated DNA Technologies (IDT) or Eurofins Genomics. The DNA and RNA products were purified via denaturing polyacrylamide gel electrophoresis (PAGE) with different percentages. Full-length products were visualized by UV shadowing and were excised from the gel. The DNA and RNA were harvested via the crush and soak method by tumbling the gel slice overnight at 4 °C in a solution of 300 mM NaCl, 10 mM Tris-HCl, and 0.1 mM EDTA (pH 7.4). Salt was removed by ethanol precipitation of the oligonucleotides twice, with two cold 70% (v/v) ethanol washes in between each precipitation. The oligonucleotides were dissolved in nuclease free water and stored at -20 °C.

### *In vitro* transcription

TracrRNA and crRNA of the *c-Myc* system were *in vitro* transcribed from T7 promoter containing template DNA by using T7 RNAP. 3 µM DNA template was transcribed in a 100 μL reaction in the presence of 1x transcription buffer (40 mM Tris-HCl, 2 mM spermidine, 10 mM DTT, and 6 mM MgCl_2_), 20 mM MgCl_2_, 2-6 mM NTPs (depending on the percentage of the individual nucleotides in the full-length transcribed RNA), and 10 μg/mL T7 RNAP. The transcribed RNAs were purified by loading them into different percentages of denaturing PAGE. UV shadowing was used to identify the full-length product band. The RNAs were extracted from the gel slice by soaking it overnight in elution buffer and then collecting the RNAs via ethanol precipitation, as previously described.

### *In vitro* RNA polymerase assay

We used an *in vitro* T7 RNA polymerase assay to investigate the effect of GQ and dCas9 on transcription. The DNA construct was prepared by annealing the template strand and non-template strand in a 1:1 molar ratio (400 nM) at 95 °C for 5-10 minutes, followed by either slow or fast cooling to room temperature. The annealing was performed at 100 mM KCl and 2 mM MgCl_2_. The gRNA components crRNAs and tracrRNA were annealed separately (each at 1.2 µM) at 95 °C for 5 minutes followed by slow cooling to room temperature. The resulting guide RNA constructs are called gR-n accordingly. Ribonucleoprotein (RNP) complexes (dCas9-gRNA) were formed by mixing the annealed gRNA constructs with dCas9 protein (Sigma-Aldrich) in 2:1 molar ratio (600 nM) in the presence of 1xCas9 buffer (20 mM Tris-HCl, 100 mM KCl, 5 mM MgCl_2_, 1 mM DTT, and 5% glycerol).

The annealed DNA construct and the RNP complex were mixed and incubated at 37 °C for 20 minutes. Following incubation, *in vitro* transcription was performed by adding all of the transcription components. During *in vitro* transcription, 2-3 aliquots were removed at different reaction times and the transcription was terminated with stop buffer [7 M urea, 10 mM Tris-HCl, and 0.1 mM EDTA (pH 7.5)]. The reaction products were separated using 8% denaturing PAGE. The gel was stained with Sybr Gold solution for 25 minutes followed by visualization on a Typhoon FLA 9500 fluorescence imager (GE Life Sciences) by selecting Cy3 scanning mode. ImageJ software was used to further process the gel image.

### Transfection of CRISPR Ribonucleoprotein (RNP)

When the differentiated Ramos cells were about 70% confluent, they were transfected with the CRISPR-dCas9 (RNP) complexes using TransIT-X2 transfection reagents (Mirus Bio, USA). TransIT-X2 reagents were stored at room temperature to warm up and gently vortexed before use in order to prepare the TransIT-X2:RNP complex for transfection. For each of the four targeting sites, gRNA (final concentration of 25 nM per well) was prepared by combining crRNA and tracrRNA at a 1:1 molar ratio, then annealing for 10 minutes at room temperature. Subsequently, the annealed product was moved into a 100 μL tube containing OptiMEM reduced serum media. Next, the precise amount of dCas9 (Sigma-Aldrich) protein stock was added to the RNP complexes, resulting in a final concentration of 12.5 nM per well and mixed gently by pipetting. For ten minutes, the RNP mixture was incubated at room temperature. The RNP mixture was pipetted with 3 μL of TransIT-X2 transfection reagent to create the TransIT-X2:RNP complex. The TransIT-X2:RNP complexes were added dropwise to various regions of the well after a 15-minute room temperature incubation period. The plate was then gently rotated to ensure that the complexes were distributed evenly. Lastly, cells treated with the TransIT-X2:RNP complex were cultured for 24 hours at 37 °C with 5% CO_2_ in a humidified incubator.

### Quantitative Reverse Transcription Polymerase Chain Reaction (RT-qPCR) Assay

To attain a more complete picture of dCas9-mediated regulation in *c-Myc* transcription, we performed a systematic cellular study in Burkitt Lymphoma Ramos cell line. Ramos cells were maintained in RPMI medium with high glucose supplemented with 20% fetal bovine serum (FBS) and 1% antibiotics (streptomycin and penicillin) at 37 °C in 5% CO_2_ in a humidified incubator. To measure *c-Myc* gene expression, Ramos cells were grown in 12-well plates at 37 °C in 5% CO_2_ in a humidified incubator for 24 h. The Ramos cells were treated with different CRISPR-dCas9 (RNP) complexes using TransIT-X2 transfection reagents (Mirus Bio, USA) when cells were ∼70% confluent. After the treatment, the cells were incubated for 24 h. The total RNA from transfected Ramos cells was extracted using the Trizol reagent following a previously optimized protocol. The complementary DNA (cDNA) was synthesized with 500 ng of total RNA using cDNA Super Mix (Quanta Biosystems, USA). The RT-qPCR assay was performed to quantify the endogenous *c-Myc* mRNA level using specific primers and SYBR Green PCR Master Mix kit (Quanta Biosystems) on an Eppendorf Mastercycler RealPlex2 Sequence Detection System. The relative fold change in *c-Myc* expression was determined using Livak Method (35).

### Western Blot

Total proteins were extracted from the Ramos cells with RIPA buffer (Santa Cruz) 24 hours after transfection with different CRISPR-dCas9 (RNP) complexes, and 50 μg of protein lysate was separated by 15% SDS−PAGE then transferred to the polyvinylidenfluoride (PVDF) membrane. Successively, they were blocked in 5% of non-fat milk in PBS + 0.1% Tween-20. Then, the blotted membranes were incubated overnight at 4 °C with primary antibodies of anti-c-Myc (1:500) (sc-40, Santa Cruz Biotechnology, Dallas, TX, USA), GAPDH was used as a loading control (G-9, sc-365062) antibody at 1:1000 dilution. Horseradish peroxidase-conjugated goat anti-mouse IgG (sc-2005) was used as secondary 86 antibody at 1:1000 dilutions. Proteins were visualized by Western Blotting Luminol Reagent (sc2048) in ChemiDoc-ItTS2 Imager.

### Cell Viability

The cell viability was determined by 3-(4,5-dimethylthiazol-2-yl)-5-(3-carboxymethoxyphenyl)-2-(4-sulfo-phenyl)-2H-tetrazolium (MTS) assay (CellTiter 96® Aqueous One Solution Cell Proliferation Assay, Promega, Madison, WI, USA). Ramos cells were seeded in 96-well plates at density of 10^5^ cells/well and incubated for 48 h prior to experimental treatments. After that, cells were treated with different CRISPR-dCas9 (RNP) complexes, 20 μL/well of MTS dye solution was added to culture medium, and cells were incubated for 8, 16, 24, 36, 48 h at 37 °C. The amount of formazan product is directly proportional to the number of living cells in culture, and it was detected by absorbance measurements at 490 nm wavelength utilizing the BioTek Synergy Neo2 Multi-detection microplate reader.

### Molecular Beacon Assay

The molecular beacon assays were performed in a BioTek Synergy Neo2 Multi-Detection Microplate Reader. For each measurement, 1 nM DNA template and 1.25 U/μL T7 RNAP (New England Biolabs) were mixed in 100 μL transcription buffer containing 40 mM Tris-HCl (pH 8.5), 50 mM KCl, 6 mM MgCl_2_, 2 mM spermidine, 1 mM dithiothreitol and 2 U inorganic pyrophosphatase. Each sample was preincubated with 500 nM molecular beacon probe (containing a fluorophore and quencher) and loaded onto a 96-well transparent plate (ThermoFisher Scientific) at 37 °C. The reaction was initiated by adding 1 mM NTP mix (New England Biolabs). Cy3 and Cy5 were excited with light at λ_ex_=540±18 nm and λ_ex_=640±20 nm, respectively. The fluorescence emission signals of Cy3 and Cy5 were collected at λ_em_=579±20 nm and λ_em_=681±20 nm, respectively. The signal was collected every 30 s. The average of the last 10 measurements (last 5 min of the 120 min experiment) were used as the saturation levels in the analysis to determine Cy5/Cy3 signal ratio.

## RESULTS

Initially, we sought to investigate the intricate relationship among the CRISPR-dCas9, target sites, PQS, template and non-template strands of a gene in terms of controlling gene expression, which guided us to create several rationally designed constructs. Fig. 1A shows a schematic of four different CRISPR-dCas9 target sites, indicated by the guide RNA strands that satisfy the PAM requirements of dCas9 (NGG in the non-target strand), in the vicinity of *c-Myc* PQS. The PQS is in the template strand (TS); therefore, targeting the sequences that overlap with the PQS in the TS is expected to destabilize the GQ. On the other hand, targeting the non-template strand (NTS) in the vicinity of PQS might stabilize GQ as it diminishes the competition with Watson-Crick pairing with the complementary strand. Specifically, gRNA-1 targets the TS and is complementary to 1^st^ and 2^nd^ G-repeats of the PQS while gRNA-2 and gRNA-3 target the NTS and overlap with sequences that are complementary to the G-repeats in the 5′and 3′ sides, respectively. The gRNA-4 targets the NTS and does not overlap with the PQS, hence should not significantly impact the GQ stability.

**Figure 1.**
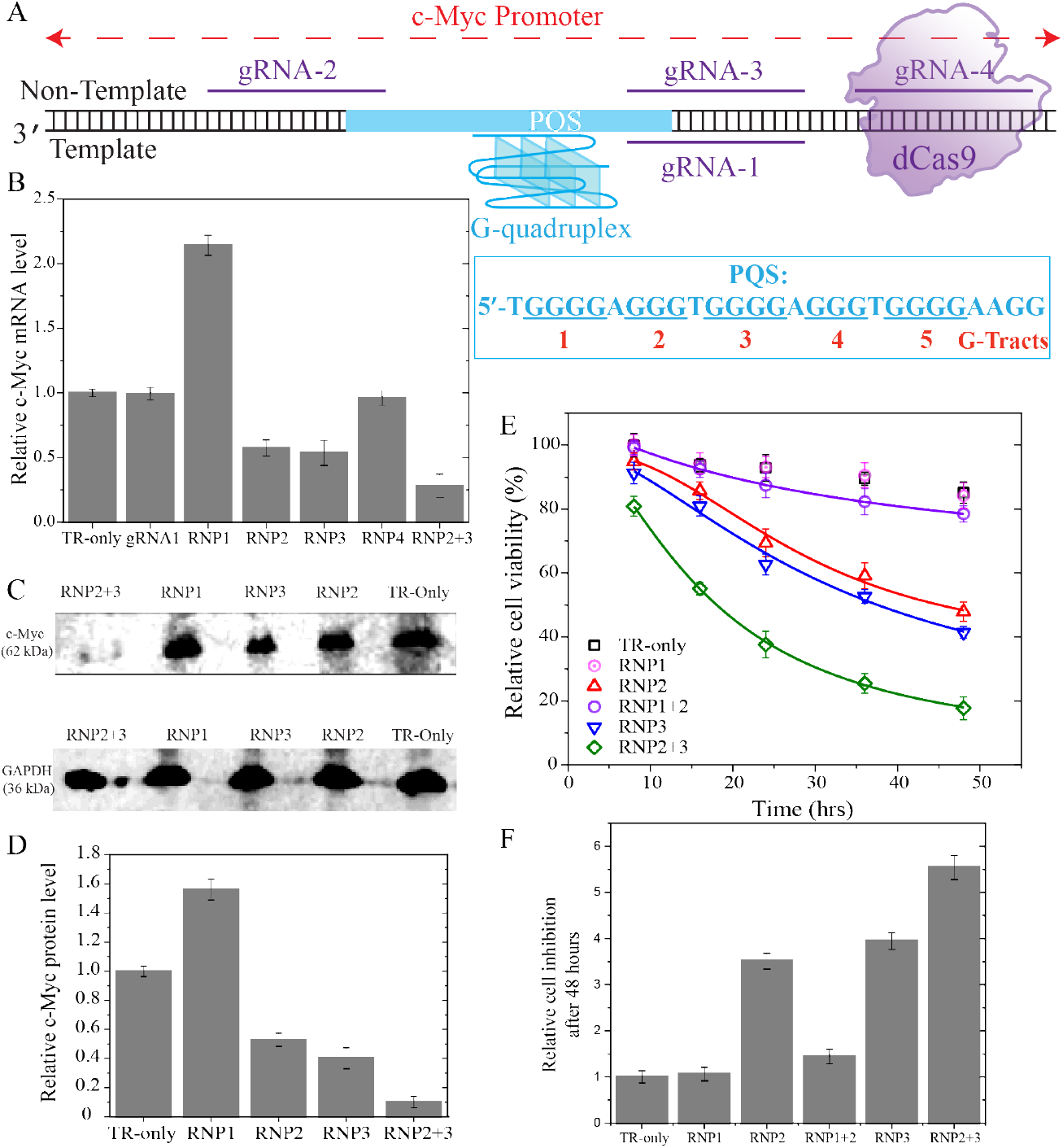
Experiments in which the vicinity of PQS was targeted with CRISPR-dCas9 in Burkitt’s lymphoma cells (Ramos cells). (A) Schematic of the DNA construct CRISPR-dCas9 target sites 1-4 on the template and non-template strands. The PQS is and the G-tracts are indicated in the schematic. (B) RT-qPCR studies illustrating suppression and elevation of c-Myc RNA levels. RNP2+3 refers to targeting sites 2 and 3 simultaneously. (C) and (D) Image of western blot and quantitation of c-Myc protein levels. (E) and (F) Cell viability studies in cells targeted with RNP1, RNP2, RNP3, RNP1+RNP2, or with RNP2+RNP3. The symbols are data points and lines are Hill function fit to the respective data. Due to small level of inhibition in cell viability in the TR-only (in the absence CRISPR-dCas9 targeting) and RNP1 cases, the data could not be fitted reliably. (F) The relative cell inhibition is quantified with respect to the TR-only case after 48 hours of introducing CRISPR-dCas9 to the cells. In (B), (C), and (F), the bars represent the average values of at least three measurements and the error bars are the standard errors associated with these measurements.

Fig. 1B-C shows results of experiments in a Burkitt’s Lymphoma cell line (Ramos cells) in which we targeted sites 1-4 with CRISPR-dCas9 (designated with RNP1-4 to highlight the ribonucleoprotein complex). The levels of *c-Myc* mRNA and protein levels after CRISPR-dCas9 treatment were quantified via RT-qPCR (Fig. 1B) and western blot (Fig. 1C), respectively. The *c-Myc* mRNA levels dropped to 0.57±0.06 and 0.57±0.09 fold of the control experiment (‘TR-Only’ where CRISPR-dCas9 was not introduced to the cells) in case of RNP2 and RNP3, respectively, while they increased by 2.14±0.07 fold in case of RNP1. In the case of RNP4, the transcription levels were similar to those of the control (0.96±0.05 fold), suggesting CRISPR-dCas9 does not present a significant blockade for transcription when this site is targeted. Similar patterns were observed for protein level suppression in case of RNP2 and RNP3 (0.53±0.04 fold and 0.40±0.07 fold, respectively). Even though significantly higher than the control (1.56±0.07 fold), the enhancement at the protein level in case of RNP1 was lower compared to the increase at mRNA level (1.56±0.07 fold in protein level compared to 2.14±0.07 at mRNA level). The statistical analyses of these data are reported in Supplementary Table S2. All cases are significantly different from the control (p<0.05) except for RNP4 case, which is not significantly different from the control.

As suppression of *c-Myc* might be of significance to inhibit cell proliferation in malignancies, we tested whether more robust suppression is achievable by targeting sites 2 and 3 simultaneously with CRISPR-dCas9. In agreement with these expectations, we observed the *c-Myc* mRNA levels dropped to 0.28 ± 0.08 of control when both RNP2 and RNP3 were introduced to the cells (Fig. 1B), which is beyond the suppression achieved with GQ-stabilizing small molecules (∼0.50-fold in the presence of 100 μM TMPyP4) (31). Even higher levels of suppression were observed at the c-Myc protein level (0.10 ± 0.04 fold of control) when both RNP2 and RNP3 were employed together (Fig. 1D). The statistical analyses of these data are reported in Supplementary Table S3. All cases are significantly different from the control (p<0.001).

We also investigated whether suppression of *c-Myc* expression impacts cell viability. Fig. 1E-F shows these studies in which the number of viable cells in multi-well plates is quantified in the absence of CRISPR-dCas9 treatment and after being treated with RNP1, RNP2, RNP3, RNP1+RNP2 (′RNP1+2′ in figures), or RNP2+RNP3 (′RNP2+3′ in figures). In case of the control sample (not treated with CRISPR-dCas9), (85±3)% of all the cells remained viable after 48 hours, while only (48 ± 3)% and (41 ± 2)% were viable in case of RNP2 and RNP3 treatment, respectively. The cells treated with RNP1, which elevates c-Myc expression, perform as well as the control, suggesting introduction of the CRISPR complex does not reduce cell viability. The fraction of viable cells was reduced to (18 ± 4)% in case of simultaneous RNP2+RNP3 treatment after 48 hours. On the other hand, cell viability was maintained at (79±3)%, which is slightly less than that of the control when RNP1 and RNP2 were introduced together. This suggests RNP1 (the enhancer) compensates for the suppression caused by RNP2. The statistical analyses of these data are reported in Supplementary Table S4. All cases are significantly different from the control (p<0.001).

Next, we investigated the mechanistic details of the suppression and activation achieved in these cellular studies using *in vitro* transcription assays. In general, RNAP transcribes RNA until elongation leads to a full-length product (Fig. 2A-B), while stalling of RNAP results in truncated RNA products (Fig. 2C-D). We designed DNA constructs in which the PQS and dCas9 target sites were placed downstream of the transcription start site (TSS) in order to probe their propensity to stall T7 RNAP progression (36). The DNA construct includes the T7 RNAP promoter, the 27-nt long c-Myc PQS (which contains five G-tracts, Fig. 1A), and the flanking sequences around the PQS to be able to use the same guide RNA oligos as those used in cellular studies (Fig. 2A). GQ formation within such a construct was confirmed with circular dichroism measurements (Supplementary Fig. S1).

**Figure 2.**
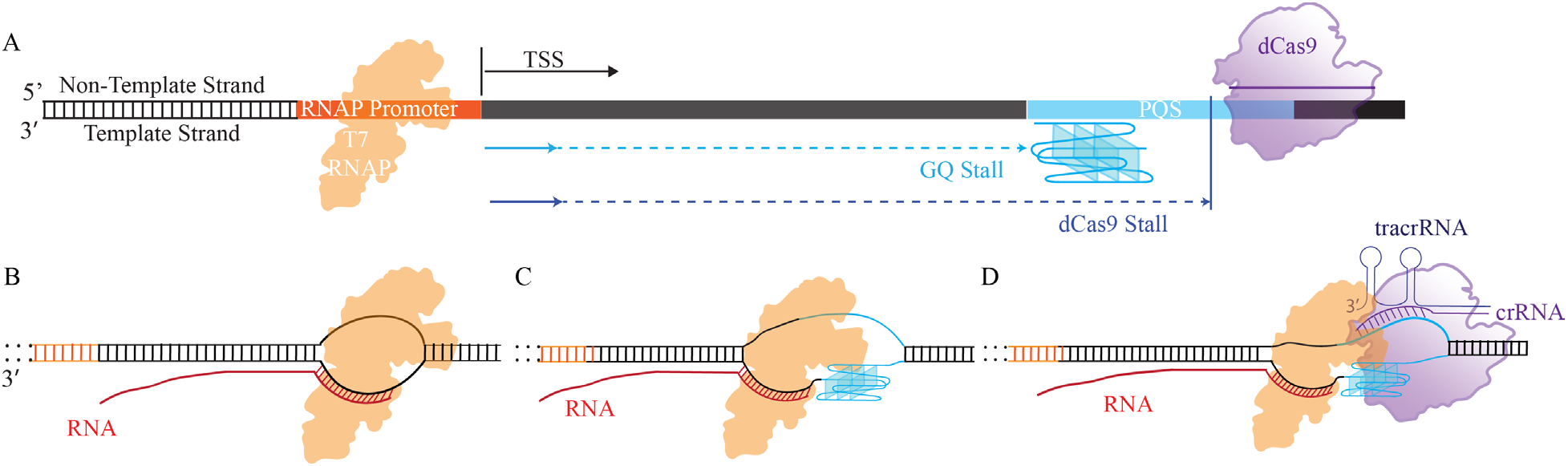
Schematic of in vitro transcription assay. (A) DNA construct indicating the relative positions of the RNAP promoter, the PQS and one of the dCas9 target sites. (B) In the absence of any blockade, RNAP completes the transcription of DNA, resulting in a full-length RNA product. (C) and (D) GQ and CRISPR-dCas9 could stall RNAP progression, resulting in truncated RNA products.

Using such constructs, we performed fluorescence-based beacon assay measurements where the products of *in vitro* transcription were quantified in real time based on the increase in fluorescence signal (37). In this assay, two short oligos that are complementary to segments of the RNA transcript either before or after the PQS were utilized (Fig. 3A-B). The beacon oligos were labeled with a Cy3 or a Cy5 fluorophore on one end and a broadband quencher at the other. When free in solution, the fluorescence signal is quenched due to flexibility of ssDNA and proximity of the dye and quencher. Upon hybridization with the complementary strand, the single-strand is stretched causing separation of the fluorophore and quencher from each other, resulting in emission of a fluorescence signal. The binding site for Cy3-labeled beacon is before the PQS and the dCas9 target sites, so we expect an increase in Cy3 signal regardless of RNAP stalls due to dCas9 or GQ. The binding site for Cy5-labeled beacon is downstream of dCas9 binding sites and the PQS, so an increase in Cy5 signal is expected only when RNAP overcomes these blockades. Therefore, the Cy3 signal correlates with the quantity of all transcription events, while the Cy5 signal correlates with the quantity of events that reach the end, meaning culmination in a full-length RNA product. The ratio of Cy5 to Cy3 signal quantitates the fraction of transcription events that by-pass the blockades, which we expect to be correlated with observed cellular transcription levels reported in Fig. 1B. In the absence of active transcription (absence of RNAP and NTP), the Cy3 and Cy5 signals due non-specific binding or binding to the non-template DNA strand was 1% (∼30 vs. ∼3000 a.u.) and 5% (∼30 vs ∼600 a.u.) of the Cy3 and Cy5 signals, respectively, observed during active transcription, suggesting >95% of all hybridization events are due to binding to transcribed RNA (Supplementary Fig. S2). The guide RNA strands used in cellular studies (gRNA-1, gRNA-2, and gRNA-3) were utilized in the beacon studies. We also designed a DNA construct (GQ-Mut) in which the GQ was significantly destabilized by mutating relevant guanines to other nucleotides. Circular dichroism measurements on the GQ-Mut construct show the absence of signature GQ motifs at 150 mM KCl (Supplementary Fig. S1). To distinguish the constructs, the unmodified construct will be referred to as the wild type (WT) construct.

**Figure 3.**
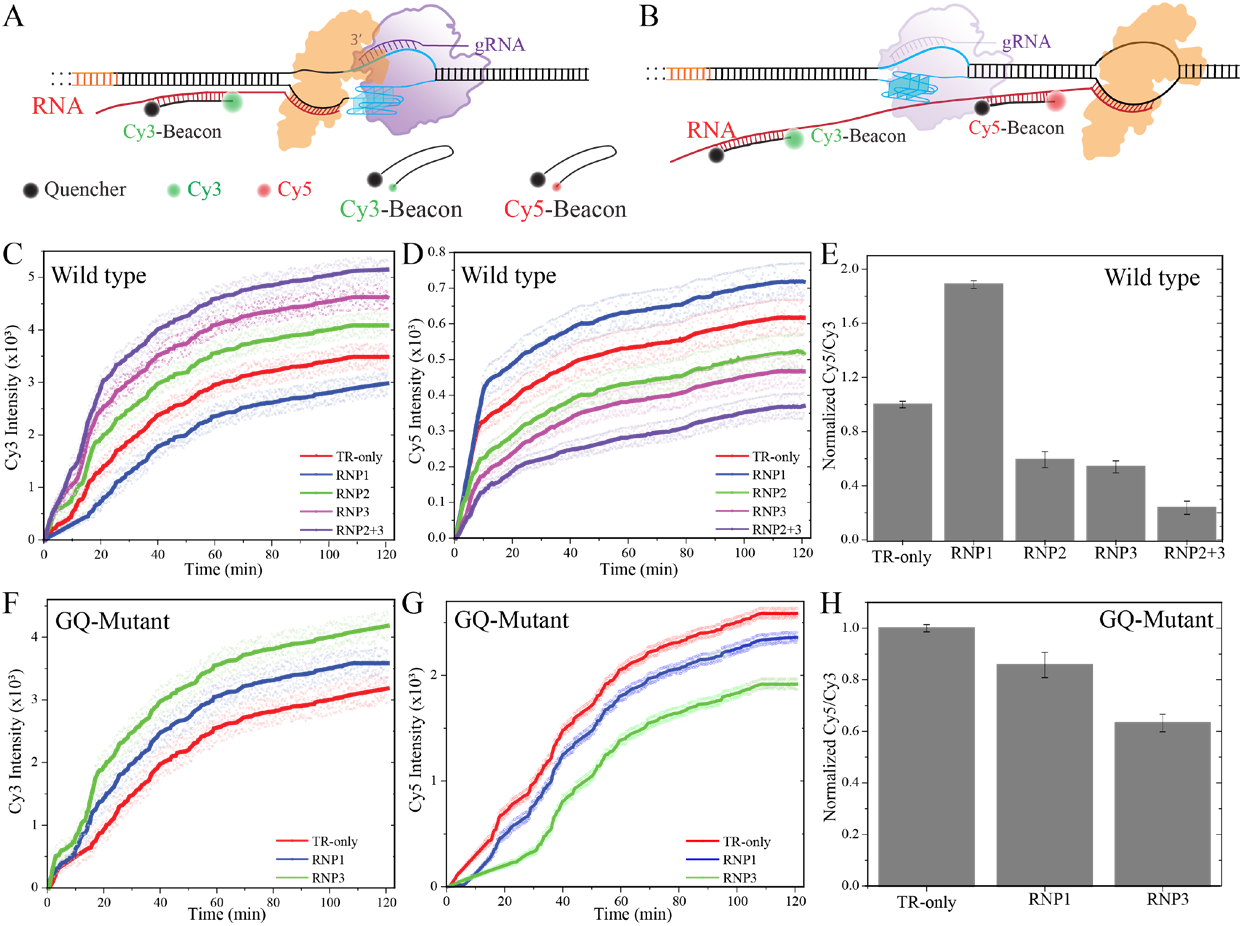
In vitro fluorescence beacon assays. (A) Schematic of the assay. The beacon strands contain a fluorophore (Cy3 or Cy5) at one end and a broad-band quencher at the other. When free in solution, the fluorescence signal is quenched due to proximity of the fluorophore and quencher. The Cy3/Cy5 beacon strand is complementary to an RNA sequence upstream/downstream of PQS and the dCas9 target sites. Binding of the beacon strands to the complementary RNA results in distancing of the fluorophore and quencher and emission of fluorescence signal. (D)-(E) and (G)-(H) Cy3 and Cy5 fluorescence emission intensities as a function of time for the wild type (contains the PQS) and GQ-mutant (PQS is mutated) DNA construct, respectively. The circles around the lines are the data points obtained from five independent measurements and the lines are the average of these measurements. (F) and (I) The ratio of Cy5/Cy3 intensities at saturation (the average of last 10 points before 120 min) for the wild type and GQ-mutant constructs, respectively. The bars are the average of five measurements and the error bars are the standard error.

Fig. 3C-D and Fig. 3F-G show the time dependent variation of the Cy3 and Cy5 fluorescence signals over a two-hour period for the WT and GQ-Mut constructs, respectively. To quantify the relative transcription events that reach full length, we plotted the Cy5/Cy3 signal ratio at saturation in Fig. 3E and Fig. 3H for the WT and the GQ-Mut constructs, respectively. Targeting the GQ-Mut construct in the NTS by RNP3 reduces the Cy5/Cy3 ratio to 0.63 ± 0.03 fold of the control (in the absence of CRISPR-dCas9) while targeting the template strand with RNP1 reduces it to 0.86 ± 0.04 fold of control. These results suggest, in the absence of a GQ structure, dCas9 presents a more prominent blockade for RNAP when the NTS is targeted compared to targeting TS. Due to mutations in the DNA sequence, RNP2 and RNP4 could not be utilized for the GQ-Mut construct.

Fig. 3C-E show similar measurements on the WT construct where RNP2 and RNP3 reduce the Cy5/Cy3 ratio to 0.59 ± 0.05 and 0.54 ± 0.04 fold of the control, respectively, while RNP1 increases it to 1.89 ± 0.02 fold of the control. Targeting the DNA by RNP2 and RNP3 simultaneously reduced the Cy5/Cy3 ratio to 0.24 ± 0.04-fold of the control. The statistical analyses of these data are reported in Supplementary Table S5 for the wild type and Table S6 for the GQ-mutant construct. All cases are significantly different from the control (p<0.01 for all cases). These results are in excellent quantitative agreement with the cellular assays presented in Fig. 1B, suggesting the Cy5/Cy3 ratio is an acceptable proxy for the observed levels of transcription regulation.

The results of these cellular and *in vitro* studies suggest suppression of *c-Myc* expression is primarily related to stalling of RNAP progression by collective action of GQ and CRISPR-dCas9 (resulting in truncated transcripts). On the other hand, enhanced transcription is primarily due to destabilization of the GQ structure (which otherwise blocks RNAP) by targeting the putative GQ forming sequence with CRISPR-dCas9. RNAP progression is more likely to be stalled by dCas9 when it is bound to the non-template strand compared to the template strand, which has been attributed to whether the collision between RNAP and dCas9 takes place in a PAM-distal (when dCas9 is bound to TS) or PAM-proximal manner (when dCas9 is bound to TS), as illustrated in Fig. 4 (38–40). Our cellular and *in vitro* beacon assay results are consistent with this model, which we further tested using an orthogonal assay that relied on the analysis of the products of an *in vitro* RNAP stop assay by gel electrophoresis. In addition to sites 1-4 of Fig. 1-3, we designed a DNA construct in which new sites (sites 5-7) that did not overlap with the PQS can be targeted in order to better identify the role of GQ in the observed transcription regulation.

**Figure 4.**
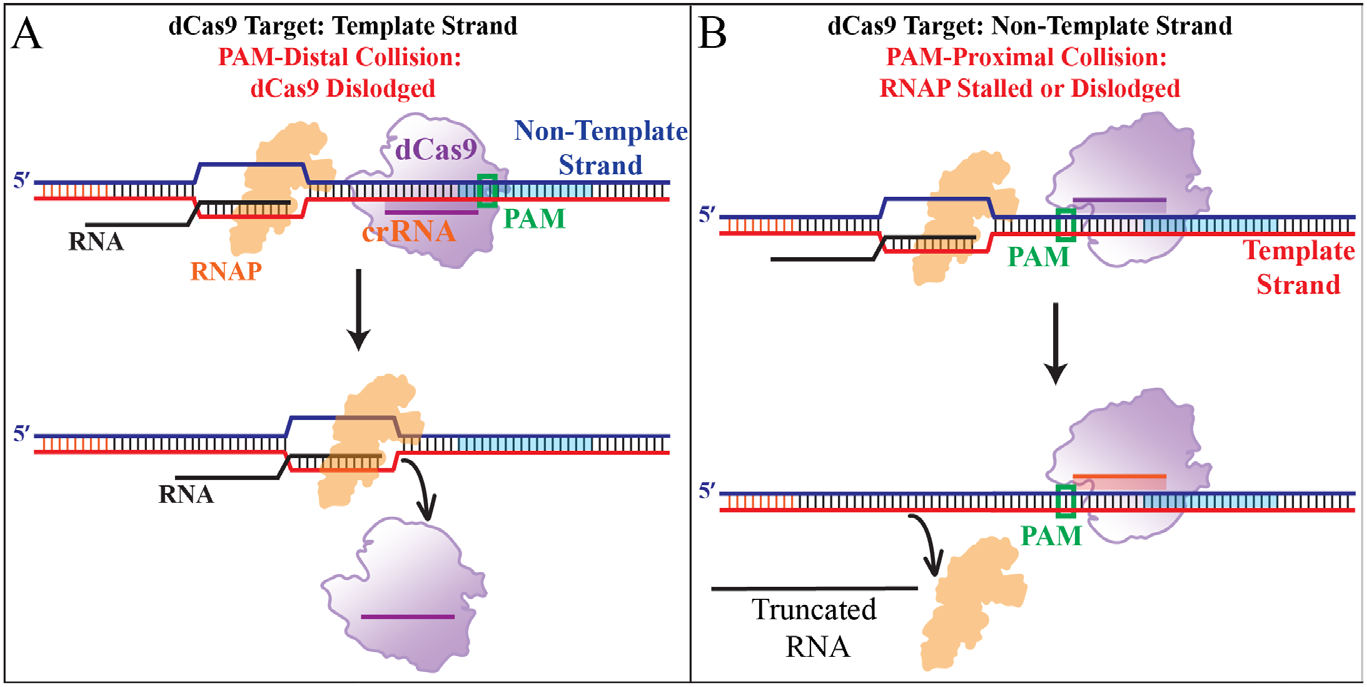
Schematic of a model representing different modes of collision between RNAP and dCas9. (A) When dCas9 targets the template strand, RNAP and dCas9 collide in a PAM-distal mode in which RNAP is more likely to dislodge dCas9 and proceed with transcription. (B) (A) When dCas9 targets the non-template strand, RNAP and dCas9 collide in a PAM-proximal mode in which RNAP is more likely to be stalled or dislodged from the DNA, both resulting in a truncated RNA transcript.

Fig. 5A shows a schematic of this DNA construct and the dCas9 target sites. Sites 1-4 are identical to those in cellular and *in vitro* beacon assays and use the same guide RNA strands (gR1-4). Sites 5-7 are upstream or downstream of PQS (without overlap) and are in either the NTS (Site 5) or TS (Sites 6-7). Sites 4-7 do not overlap with the PQS; hence, targeting them with dCas9 should not significantly impact the GQ stability.

**Figure 5.**
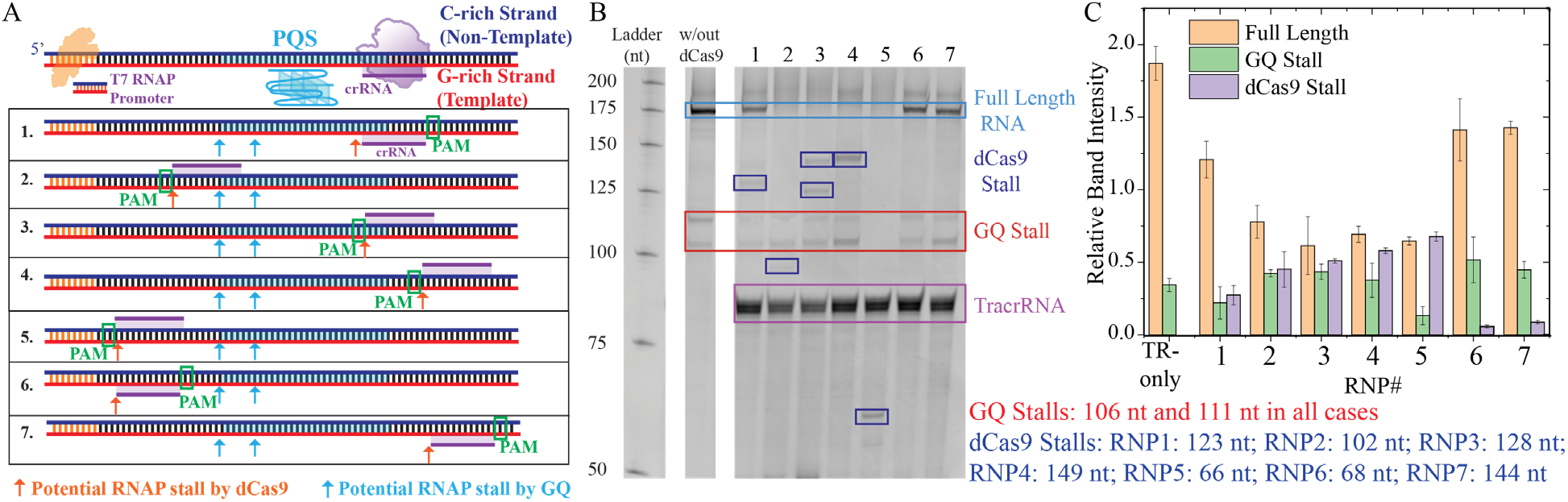
Gel electrophoresis measurements analyzing the products of an in vitro transcription assay (RNAP stop assay). (A) Schematic of the DNA construct that includes a T7 RNAP promoter, PQS and dCas9 target sites. The PQS and dCas9 target sites are downstream of RNAP promoter; therefore, stalling of the RNAP at these sites results in truncated RNA transcripts. In the schematic below, the seven dCas9 target sites are indicated, in addition to GQ stall sites (cyan arrows) and dCas9 stall sites (red arrows). The expected RNA transcripts corresponding to these stalls are listed on the right (under (C)). (B) Gel image showing the products of the in vitro transcription assay. GQ stalls are indicated with a red rectangle and dCas9 stalls are indicated with dark blue rectangles. (C) Quantitation of stall bands for the control sample (TR-only in which CRISPR-dCas9 was excluded from the assay) and the seven CRISPR-dCas9 target sites (RNP1-7). The bars represent averages of at least three measurements and the error bars are the standard error of these measurements.

The potential stall sites for RNP1-7 complexes and those for GQ have been marked in the schematic in Fig. 5A and on the gel electrophoresis image in Fig. 5B (the expected lengths of the truncated RNA products are given in Fig. 5). The five G-tracts in the PQS are adequate for formation of at least two different GQs which result in two stall bands for all cases (marked with one blue rectangle as they are all of the same length), except RNP5. The stall products due to CRISPR-dCas9 complexes are marked with red boxes. We observed dCas9 stalls in all four cases (RNP2-5) when dCas9 targeted the non-template strand. On the other hand, we observed dCas9 stall in only one of the three cases (RNP1 but not RNP6 or RNP7) when dCas9 targeted the template strand. The band intensities for these stalls are quantified in Fig. 5C. The statistical analyses of these data are reported in Supplementary Table S7. All cases are significantly different from the control (p<0.01 for all cases).

These results also suggest dCas9 induced RNAP stalls are more prominent when the non-template strand is targeted. This prominent stalling of RNAP by dCas9 when NTS is targeted might explain why a GQ stall band is not observed in case of RNP5 where the RNAP is stalled by the dCas9 (bound to the NTS) before it reaches the PQS.

## DISCUSSION

We show the synergistic roles CRISPR-dCas9 and G-quadruplex structures can play in modulating the *c-Myc* expression at RNA and protein levels in Burkitt’s lymphoma cells. We also show that cell viability is inhibited at a level correlated with the observed suppression in *c-Myc* levels. In addition, we report extensive measurements using *in vitro* transcription and polymerase stop assays and deduce mechanistic insights into the underlying processes. We observe elevated levels of truncated products when dCas9 targeted the non-template strand compared to targeting the template strand. While this feature alone yielded varying levels of transcription suppression in the absence of PQS, we showed that it is possible to attain elevated transcription and a broader range of suppression when the PQS is targeted. GQ structures, when in TS, as is the case in *c-Myc*, can block RNAP progression (41, 42), which serves as the second critical element of the observed modulation. Targeting the NTS with dCas9 stabilizes the GQ structure, which further strengthens the blockade effect and suppression of transcription. On the other hand, targeting the sequences in TS that at least partially overlap with the PQS, destabilizes the GQ. In this case, whether the blockade is stronger or weaker depends on whether the reduced stabilization of the GQ is as significant as the blockade presented by binding of CRISPR-dCas9 to TS. In the case of *c-Myc*, we illustrate an enhanced transcription when the vicinity of PQS in the TS is targeted, which suggests the reduced stability of the GQ is the dominant effect. Measurements in control constructs that lack the PQS show that CRISPR-dCas9 does not present a significant blockade for the RNAP when it targets the template strand, which further supports the overall picture that emerges from these studies. As *c-Myc* is critical for cell proliferation and is upregulated in diverse cancers, effective suppression of its expression is desirable. We demonstrated that this can be achieved by targeting the NTS at two sites simultaneously, on the 3′ and 5′ sides of the complementary sequence to PQS. The level of suppression attained in this case at the RNA and protein levels is 2 to 5 fold greater than what has been reported with drug-like small molecules, which lack sequence specificity. This effective suppression is also reflected in cell viability assays in which we observe significantly higher inhibition when these two sites are simultaneously targeted with dCas9. We also observe a complementary effect when RNP1 (enhancer) and RNP2 (suppressor) are both introduced to the cells. In this case, RNP1-induced enhancement compensates for RNP2-induced suppression and cell viability is minimally impacted.

Our study illustrates the potential of using CRISPR-dCas9 to target PQS that are present in numerous regulatory sites for transcription regulation. The G-rich nature of GQ forming sequences and that of PAM sequence of dCas9 (‘5′-NGG’ in the non-target strand) provide many targetable sites in the non-template strand (for systems in which the PQS is in the template strand as in *c-Myc*). This creates an ideal setting for not only transcription suppression, but also the feasibility of transcription activation (without the need for fusing dCas9 with activator proteins) when the template strand is targeted, as we demonstrated for *c-Myc*. Similar mechanisms might be valid for other Cas proteins (such as Cas12 which has a T-rich PAM), which would broaden the range of potential target sites and enable higher levels of control on gene expression. Our study illustrates an example of this great potential at RNA, protein, and cellular levels in a medically significant system.

## SUPPLEMENTARY INFORMATION

Supplementary Information is available online.

## References

1. Horvath, P. and Barrangou, R. (2010) CRISPR/Cas, the Immune System of Bacteria and Archaea. Science (1979), 327, 167–170.

2. Barrangou, R., Fremaux, C., Deveau, H., Richards, M., Boyaval, P., Moineau, S., Romero, D.A. and Horvath, P. (2007) CRISPR Provides Acquired Resistance Against Viruses in Prokaryotes. Science (1979), 315, 1709–1712.

3. Qi, L.S., Larson, M.H., Gilbert, L.A., Doudna, J.A., Weissman, J.S., Arkin, A.P. and Lim, W.A. (2013) Repurposing CRISPR as an RNA-Guided Platform for Sequence-Specific Control of Gene Expression. Cell, 152, 1173–1183.

4. Larson, M.H., Gilbert, L.A., Wang, X., Lim, W.A., Weissman, J.S. and Qi, L.S. (2013) CRISPR interference (CRISPRi) for sequence-specific control of gene expression. Nat Protoc, 8, 2180–2196.

5. Konermann, S., Brigham, M.D., Trevino, A.E., Joung, J., Abudayyeh, O.O., Barcena, C., Hsu, P.D., Habib, N., Gootenberg, J.S., Nishimasu, H., et al. (2015) Genome-scale transcriptional activation by an engineered CRISPR-Cas9 complex. Nature, 517, 583–588.

6. Bikard, D., Jiang, W., Samai, P., Hochschild, A., Zhang, F. and Marraffini, L.A. (2013) Programmable repression and activation of bacterial gene expression using an engineered CRISPR-Cas system. Nucleic Acids Res, 41, 7429–7437.

7. Gilbert, L.A., Horlbeck, M.A., Adamson, B., Villalta, J.E., Chen, Y., Whitehead, E.H., Guimaraes, C., Panning, B., Ploegh, H.L., Bassik, M.C., et al. (2014) Genome-Scale CRISPR-Mediated Control of Gene Repression and Activation. Cell, 159, 647–661.

8. Chavez, A., Scheiman, J., Vora, S., Pruitt, B.W., Tuttle, M., P R Iyer, E., Lin, S., Kiani, S., Guzman, C.D., Wiegand, D.J., et al. (2015) Highly efficient Cas9-mediated transcriptional programming. Nat Methods, 12, 326–328.

9. Chen, S., Sanjana, N.E., Zheng, K., Shalem, O., Lee, K., Shi, X., Scott, D.A., Song, J., Pan, J.Q., Weissleder, R., et al. (2015) Genome-wide CRISPR Screen in a Mouse Model of Tumor Growth and Metastasis. Cell, 160, 1246–1260.

10. Dominguez, A.A., Lim, W.A. and Qi, L.S. (2016) Beyond editing: repurposing CRISPR–Cas9 for precision genome regulation and interrogation. Nat Rev Mol Cell Biol, 17, 5–15.

11. He, M., Zhou, X., Li, Z., Yin, X., Han, W., Zhou, J., Sun, X., Liu, X., Yao, D. and Liang, H. (2022) Programmable Transcriptional Modulation with a Structured RNA-Mediated CRISPR-dCas9 Complex. J Am Chem Soc, 144, 12690–12697.

12. Widom, J.R., Rai, V., Rohlman, C.E. and Walter, N.G. (2019) Versatile transcription control based on reversible dCas9 binding. RNA, 25, 1457–1469.

13. Anderson, D.A. and Voigt, C.A. (2021) Competitive dCas9 binding as a mechanism for transcriptional control. Mol Syst Biol, 17, e10512.

14. Bikard, D., Jiang, W., Samai, P., Hochschild, A., Zhang, F. and Marraffini, L.A. (2013) Programmable repression and activation of bacterial gene expression using an engineered CRISPR-Cas system. Nucleic Acids Res, 41, 7429–7437.

15. Larson, M.H., Gilbert, L.A., Wang, X., Lim, W.A., Weissman, J.S. and Qi, L.S. (2013) CRISPR interference (CRISPRi) for sequence-specific control of gene expression. Nat Protoc, 8, 2180–2196.

16. Qi, L.S., Larson, M.H., Gilbert, L.A., Doudna, J.A., Weissman, J.S., Arkin, A.P. and Lim, W.A. (2013) Repurposing CRISPR as an RNA-Guided Platform for Sequence-Specific Control of Gene Expression. Cell, 152, 1173–1183.

17. Hoque, M.E., Mustafa, G., Basu, S. and Balci, H. (2021) Encounters between Cas9/dCas9 and G-Quadruplexes: Implications for Transcription Regulation and Cas9-Mediated DNA Cleavage. ACS Synth Biol, 10.1021/acssynbio.1c00067.

18. Maizels, N. and Gray, L.T. (2013) The G4 genome. PLoS Genet, 9, e1003468.

19. Varshney, D., Spiegel, J., Zyner, K., Tannahill, D. and Balasubramanian, S. (2020) The regulation and functions of DNA and RNA G-quadruplexes. Nat Rev Mol Cell Biol, 21, 459–474.

20. Hänsel-Hertsch, R., Di Antonio, M. and Balasubramanian, S. (2017) DNA G-quadruplexes in the human genome: Detection, functions and therapeutic potential. Nat Rev Mol Cell Biol, 18, 279–284.

21. Marsico, G., Chambers, V.S., Sahakyan, A.B., McCauley, P., Boutell, J.M., Antonio, M.Di and Balasubramanian, S. (2019) Whole genome experimental maps of DNA G-quadruplexes in multiple species. Nucleic Acids Res, 47, 3862–3874.

22. Huppert, J.L. and Balasubramanian, S. (2007) G-quadruplexes in promoters throughout the human genome. Nucleic Acids Res, 35, 406–413.

23. Eddy, J. and Maizels, N. (2006) Gene function correlates with potential for G4 DNA formation in the human genome. Nucleic Acids Res, 34, 3887–3896.

24. Marcu, K.B., Bossone, S.A. and Patel, A.J. (2003) myc FUNCTION AND REGULATION. 10.1146/annurev.bi.61.070192.004113, 61, 809–858.

25. Pelengaris, S., Rudolph, B. and Littlewood, T. (2000) Action of Myc in vivo — proliferation and apoptosis. Curr Opin Genet Dev, 10, 100–105.

26. Thompson, E.B. (1998) The many roles of c-myc in apoptosis. Annu Rev Physiol, 60, 575–600.

27. Gabay, M., Li, Y. and Felsher, D.W. (2014) MYC activation is a hallmark of cancer initiation and maintenance. Cold Spring Harb Perspect Med, 4.

28. Dhanasekaran, R., Deutzmann, A., Mahauad-Fernandez, W.D., Hansen, A.S., Gouw, A.M. and Felsher, D.W. (2022) The MYC oncogene - the grand orchestrator of cancer growth and immune evasion. Nat Rev Clin Oncol, 19, 23–36.

29. Postel, E.H., Mango, S.E. and Flint, S.J. (1989) A Nuclease-Hypersensitive Element of the Human c-myc Promoter Interacts with a Transcription Initiation Factor. Mol Cell Biol, 9, 5123–5133.

30. Sakatsume, O., Tsutsui, H., Wang, Y., Gao, H., Tang, X., Yamauchi, T., Murata, T., Itakura, K. and Yokoyama, K.K. (1996) Binding of THZif-1, a MAZ-like Zinc Finger Protein to the Nuclease-hypersensitive Element in the Promoter Region of the c-MYC Protooncogene*. J Biol Chem, 271, 31322–31333.

31. Siddiqui-Jain, A., Grand, C.L., Bearss, D.J. and Hurley, L.H. (2002) Direct evidence for a G-quadruplex in a promoter region and its targeting with a small molecule to repress c-MYC transcription. Proc Natl Acad Sci U S A, 99, 11593–11598.

32. Esain-Garcia, I., Kirchner, A., Melidis, L., de Cesaris Araujo Tavares, R., Dhir, S., Simeone, A., Yu, Z., Madden, S.K., Hermann, R., Tannahill, D., et al. (2024) G-quadruplex DNA structure is a positive regulator of MYC transcription. Proc Natl Acad Sci U S A, 121, e2320240121.

33. Balci, H., Globyte, V. and Joo, C. (2021) Targeting G-quadruplex Forming Sequences with Cas9. ACS Chem Biol, 16, 596–603.

34. Siddiqui-Jain, A., Grand, C.L., Bearss, D.J. and Hurley, L.H. (2002) Direct evidence for a G-quadruplex in a promoter region and its targeting with a small molecule to repress c-MYC transcription. Proceedings of the National Academy of Sciences, 99, 11593–11598.

35. Livak, K.J. and Schmittgen, T.D. (2001) Analysis of relative gene expression data using real-time quantitative PCR and the 2(-Delta Delta C(T)) Method. Methods, 25, 402–408.

36. Farhath, M.M., Thompson, M., Ray, S., Sewell, A., Balci, H. and Basu, S. (2015) G-Quadruplex-Enabling Sequence within the Human Tyrosine Hydroxylase Promoter Differentially Regulates Transcription. Biochemistry, 54, 5533–5545.

37. Lee, C.Y., Joshi, M., Wang, A. and Myong, S. (2024) 5’UTR G-quadruplex structure enhances translation in size dependent manner. Nat Commun, 15.

38. Clarke, R., Heler, R., MacDougall, M.S., Yeo, N.C., Chavez, A., Regan, M., Hanakahi, L., Church, G.M., Marraffini, L.A. and Merrill, B.J. (2018) Enhanced Bacterial Immunity and Mammalian Genome Editing via RNA-Polymerase-Mediated Dislodging of Cas9 from Double-Strand DNA Breaks. Mol Cell, 71, 42-55.e8.

39. Hall, P.M., Inman, J.T., Fulbright, R.M., Le, T.T., Brewer, J.J., Lambert, G., Darst, S.A. and Wang, M.D. (2022) Polarity of the CRISPR roadblock to transcription. Nature Structural & Molecular Biology |, 29, 1217–1227.

40. Vigouroux, A., Oldewurtel, E., Cui, L., Bikard, D. and van Teeffelen, S. (2018) Tuning dCas9’s ability to block transcription enables robust, noiseless knockdown of bacterial genes. Mol Syst Biol, 14.

41. Lee, C.Y., McNerney, C., Ma, K., Zhao, W., Wang, A. and Myong, S. (2020) R-loop induced G-quadruplex in non-template promotes transcription by successive R-loop formation. Nat Commun, 11.

42. Hwang, J., Lee, C.-Y., Paul, T., Lee, H., Ha, T. and Myong, S. (2024) DNA supercoiling-mediated G4/R-loop formation tunes transcription by controlling the access of RNA polymerase. Biophys J, 123.

