## Supplementary Information for "Combining the CRISPR Activation and Interference Capabilities Using dCas9 and G-Quadruplex Structures"

### Sequences of Nucleic Acids

Table S1 lists the sequences of the strands used throughout the manuscript, with reference to the specific figures they were employed. The same TracrRNA was used to construct all the guide RNA molecules, while the CRISPR-RNA (crRNA) strands changed, i.e. gRNA-1 refers to the combination of TracrRNA and crRNA-1. The c-Myc putative G-quadruplex sequence (PQS) is marked with cyan fonts in the template strand and the corresponding mutations to eliminate GQ are marked with red fonts in the template strand of GQ-Mutant construct.

**Table S1:** *Oligonucleotide sequences used for in vitro and in vivo assays*

| Name | Sequence (5' to 3') | Figure |
|---|---|---|
| **Non-Template Strand (C-rich)** | GCTAATACGACTCACTATAGGAGCAAAAGAAAATGGTAGGCGCGCGTA GTTAATTCATGCGGCTCTCTTACTCTGTTTACATCCTAGAGCTAGAGTGC TCGGCTGCCCGGCTGAGTCTCCTCCCCACCTTCCCCACCCTCCCCACCC TCCCCATAAGCGCCCCTCCCGGGTTCCCAAAGCAGAGGGCGTGGGGG | Fig. 2, 3, 5 |
| **Template Strand (G-rich)** | CCCCCACGCCCTCTGCTTTGGGAACCCGGGAGGGGCGCTTATGGGGAG GGTGGGGAGGGTGGGGAAGGTGGGGAGGAGACTCAGCCGGGCAGCCG AGCACTCTAGCTCTAGGATGTAAACAGAGTAAGAGAGCCGCATGAATTA ACTACGCGCGCCTACCATTTTCTTTTGCTCCTATAGTGAGTCGTATTAGC | Fig. 2, 3, 5 |
| **TracrRNA** | GGAACCAUUCAAAACAGCAUAGCAAGUUAAAAUAAGGCUAGUCCGU UAUCAACUUGAAAAAGUGGCACCGAGUCGGUGCUUUUUUU | Fig. 1, 3, 5 |
| **crRNA-1** | GGCTCCCCATAAGCGCCCCTCCGUUUUAGAGCUAUGCUGUUUUG | Fig. 1, 3, 5 |
| **crRNA-2** | GGAGGGTGGGGAAGGTGGGGGUUUUAGAGCUAUGCUGUUUUG | Fig. 1, 3, 5 |
| **crRNA-3** | GGACCCGGGAGGGGCGCTTATGGUUUAGAGCUAUGCUGUUUG | Fig. 1, 3, 5 |
| **crRNA-4** | GGCAGCCGAGCACTCTAGCTCTGUUUUAGAGCUAUGCUGUUUUG | Fig. 1, 5 |
| **crRNA-5** | GGCGCCCTCTGCTTTGGGAACCGUUUUAGAGCUAUGCUGUUUUG | Fig. 5 |
| **crRNA-6** | GGAGCTAGAGTGCTCGGCTGCCGUUUUAGAGCUAUGCUGUUUUG | Fig. 5 |
| **crRNA-7** | GGTCCCGGGTTCCCAAAGCAGAGUUUUAGAGCUAUGCUGUUUUG | Fig. 5 |
| **Cy3-Beacon** | /5Cy3/CCGCATGAA/TAO/TTAACTACG/Quencher/ | Fig. 2 |
| **Cy5-Beacon** | /5Cy5/CGCCCTCTGCTTTGGGAA/3IAbRQSp/ | Fig. 2 |
| **GAPDH-Forward** | AGGTCGGTGTGAACGGATTTG | Fig. 1C-D |
| **GAPDH-Reverse** | GGGGTCGTTGATGGCAACA | Fig. 1C-D |
| **c-Myc Forward** | CCTGGTGCTCCATGAGGAGAC | Fig. 1B-D |
| **c-Myc Reverse** | CAGACTCTGACCTTTTGCCAGG | Fig. 1B-D |
| **Template Strand: GQ Mutant** | CCCCCACGCCCTCTGCTTTGGGAACCCGGGAGGGGCGCTTATGGGGAG TGTGGAGAGTGTGAGGAAGGTGGGGAGGAGACTCAGCCGGGCAGCCG AGCACTCTAGCTCTAGGATGTAAACAGAGTAAGAGAGCCGCATGAATTA ACTACGCGCGCCTACCATTTTCTTTTGCTCCTATAGTGAGTCGTATTAGC | Fig. 3G-I |
| **Non-Template Strand: GQ-Mutant** | GCTAATACGACTCACTATAGGAGCAAAAGAAAATGGTAGGCGCGCGTA GTTAATTCATGCGGCTCTCTTACTCTGTTTACATCCTAGAGCTAGAGTGC TCGGCTGCCCGGCTGAGTCTCCTCCCCACCTTCCTCACACTCTCCACAC TCCCCATAAGCGCCCCTCCCGGGTTCCCAAAGCAGAGGGCGTGGGGG | Fig. 3G-I |

### Statistical Analysis

Tables S2-S7 present the T-test analysis we performed on all the data presented in the figures to establish the significance of the observed differences under different conditions.

**Table S2:** T-test analysis on the data presented in Figure 1B.

| Name | T- Test | Comment |
|---|---|---|
| gRNA1 | t (2) = 0.23, p=0.8421 | No Significant difference |
| RNP1 | t (2) = -14.87, p<0.001 | Significantly different |
| RNP2 | t (2) =6.42, p=0.003 | Significantly different |
| RNP3 | t (2) =4.86, p=0.0082 | Significantly different |
| RNP4 | t (2) =0.68, p=0.5401 | No Significant difference |
| RNP2+3 | t (2) =8.42, p=0.0011 | Significantly different |

**Table S3:** T-test analysis on the data presented in Figure 1D.

| Name | T- Test | Comment |
|---|---|---|
| RNP1 | t (2) =-12.25, p<0.001 | Significantly different |
| RNP2 | t (2) =14.33, p<0.001 | Significantly different |
| RNP3 | t (2) =12.95, p<0.001 | Significantly different |
| RNP2+3 | t (2) =29.26, p<0.001 | Significantly different |

**Table S4:** T-test analysis on the data presented in Figure 1F.

| Name | T- Test | Comment |
|---|---|---|
| RNP1 | t (2) = -0.471, p=0.6621 | No Significant difference |
| RNP1+2 | t (2) = -3.106, p=0.038 | Significantly different |
| RNP2 | t (2) =-20.34, p<0.001 | Significantly different |
| RNP3 | t (2) =-22.92, p<0.001 | Significantly different |
| RNP2+3 | t (2) =-27.06, p<0.001 | Significantly different |

**Table S5:** T-test analysis on the data presented in Figure 3E.

| Name | T- Test | Comment |
|---|---|---|
| RNP1 | t (4) =-54.7, p<0.001 | Significantly different |
| RNP2 | t (4) =13.17, p<0.001 | Significantly different |
| RNP3 | t (4) =17.84, p<0.001 | Significantly different |
| RNP2+3 | t (4) =29.42, p<0.001 | Significantly different |

**Table S6:** T-test analysis on the data presented in Figure 3H.

| Name | T- Test | Comment |
|---|---|---|
| RNP1 | t (4) =5.86, p=0.0043 | Significantly different |
| RNP3 | t (4) =20.25, p<0.001 | Significantly different |

**Table S7:** T-test analysis on the data presented in Figure 5.

| **Name** | **T- Test** | **Comment** |
|---|---|---|
| RNP1: Full Length | t (2) =7.03, p=0.0023 | Significantly different |
| RNP1: GQ Stall | t (2) =18.39, p<0.001 | Significantly different |
| RNP1: dCas9 Stall | t (2) =22.13, p<0.001 | Significantly different |
| RNP2: Full Length | t (2) =12.13, p<0.001 | Significantly different |
| RNP2: GQ Stall | t (2) =22.47, p<0.001 | Significantly different |
| RNP2: dCas9 Stall | t (2) =15.14, p<0.001 | Significantly different |
| RNP3: Full Length | t (2) =9.88, p<0.001 | Significantly different |
| RNP3: GQ Stall | t (2) =20.49, p<0.001 | Significantly different |
| RNP3: dCas9 Stall | t (2) =21.36, p<0.001 | Significantly different |
| RNP4: Full Length | t (2) =16.93, p<0.001 | Significantly different |
| RNP4: GQ Stall | t (2) =16.56, p<0.001 | Significantly different |
| RNP4: dCas9 Stall | t (2) =20.27, p<0.001 | Significantly different |
| RNP5: Full Length | t (2) =18.89, p<0.001 | Significantly different |
| RNP5: GQ Stall | t (2) =24.08, p<0.001 | Significantly different |
| RNP5: dCas9 Stall | t (2) =18.06, p<0.001 | Significantly different |
| RNP6: Full Length | t (2) =3.38, p<0.001 | Significantly different |
| RNP6: GQ Stall | t (2) =12.59, p<0.001 | Significantly different |
| RNP6: dCas9 Stall | t (2) =28.39, p<0.001 | Significantly different |
| RNP7: Full Length | t (2) =6.53, p=0.0028 | Significantly different |
| RNP7: GQ Stall | t (2) =20.38, p<0.001 | Significantly different |
| RNP7: dCas9 Stall | t (2) =27.93, p<0.001 | Significantly different |

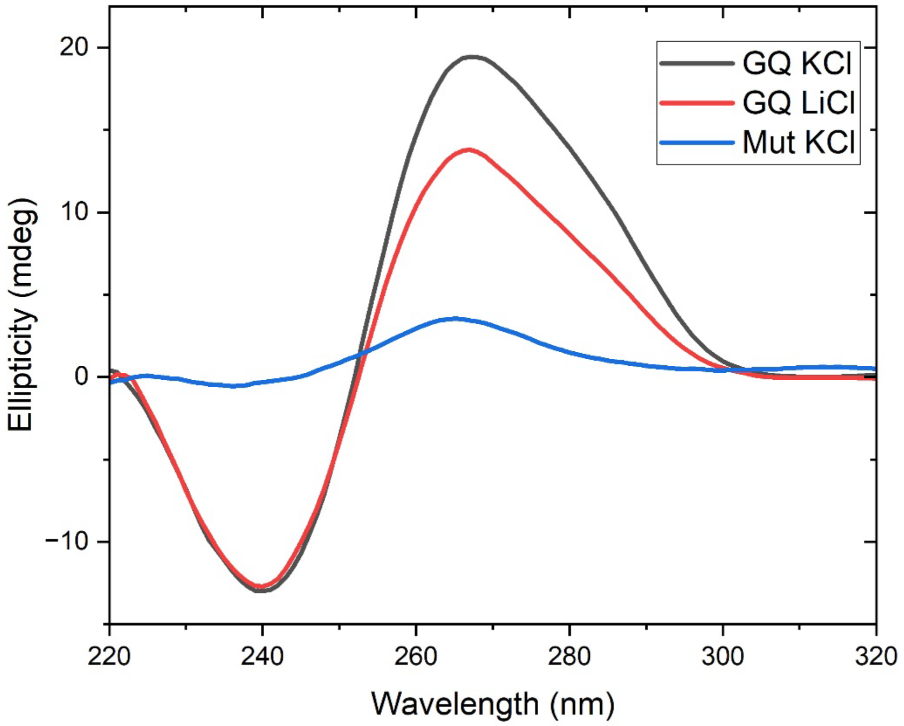


**Figure S1.** Circular dichroism spectra of the c-Myc DNA construct (the wild type) in 100 mM KCl or 100 mM LiCl. We also show the CD spectrum of GQ-Mut construct in 150 mM KCl. The DNA constructs were created by annealing the Template and Non-Template strands given in Table S1 (two first two entries for the wild type and the last two entries for the GQ-mutant construct). The wild type construct was used in Fig. 3 and Fig. 5 while the GQ-Mut construct was used in Fig. 3. The peak at 265 nm and trough at 240 nm are consistent with parallel conformation. The more prominent peak in KCl suggests a more prominent GQ formation in KCl compared to LiCl. Compared to the wild type construct, the GQ-Mut construct shows a much weaker signal suggesting the absence of the GQ.

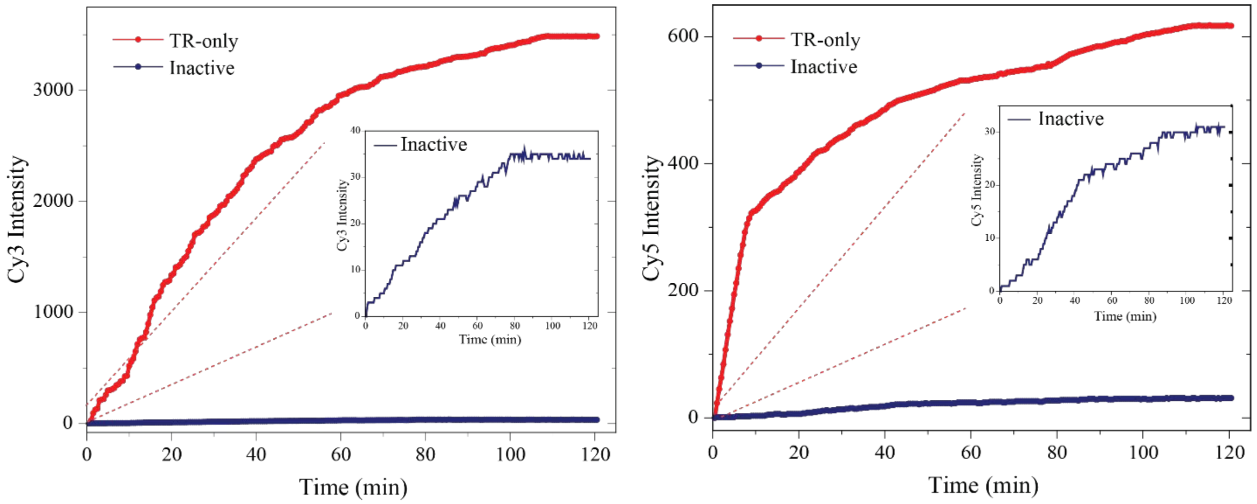


**Figure S2.** Beacon assay measurements in the absence of active transcription ("Inactive" due to absence of RNAP and NTP) or when active transcription was ongoing ("TR-Only" as described in manuscript). The Cy3 signal in the Inactive case is ~1% of that of the active case (~35 a.u. vs ~3500 a.u. at saturation). The Cy5 signal in the Inactive case is ~5% of that of the active case (~30 a.u. vs ~600 a.u. at saturation). This assay establishes the level of beacon intensity due to non-specific binding or due to binding to the non-template strand of DNA to 1-5% of that of binding to RNA molecules.
